# Live cell imaging of the hyperthermophilic archaeon *Sulfolobus acidocaldarius* identifies complementary roles for two ESCRTIII homologues in ensuring a robust and symmetric cell division

**DOI:** 10.1101/2020.02.18.953042

**Authors:** Andre Arashiro Pulschen, Delyan R. Mutavchiev, Kim Nadine Sebastian, Jacques Roubinet, Marc Roubinet, Gabriel Tarrason Risa, Marleen van Wolferen, Chantal Roubinet, Siân Culley, Gautam Dey, Sonja-Verena Albers, Ricardo Henriques, Buzz Baum

## Abstract

Live-cell imaging has revolutionized our understanding of dynamic cellular processes in bacteria and eukaryotes. While similar techniques have recently been applied to the study of halophilic archaea, our ability to explore the cell biology of thermophilic archaea is limited, due to the technical challenges of imaging at high temperatures. Here, we report the construction of the *Sulfoscope*, a heated chamber that enables live-cell imaging on an inverted fluorescent microscope. Using this system combined with thermostable fluorescent probes, we were able to image *Sulfolobus* cells as they divide, revealing a tight coupling between changes in DNA compaction, segregation and cytokinesis. By imaging deletion mutants, we observe important differences in the function of the two ESCRTIII proteins recently implicated in cytokinesis. The loss of CdvB1 compromises cell division, causing occasional division failures and fusion of the two daughter cells, whereas the deletion of *cdvB2* leads to a profound loss of division symmetry, generating daughter cells that vary widely in size and eventually generating ghost cells. These data indicate that DNA separation and cytokinesis are coordinated in *Sulfolobus*, as is the case in eukaryotes, and that two contractile ESCRTIII polymers perform distinct roles to ensure that *Sulfolobus* cells undergo a robust and symmetrical division. Taken together, the *Sulfoscope* has shown to provide a controlled high temperature environment, in which cell biology of *Sulfolobus* can be studied in unprecedent details.

## Introduction

Live imaging is essential for the understanding of complex dynamic cellular events, such as changes in cell shape, cell division and DNA segregation. Despite such methods being well-established in bacteria and eukaryotes, live imaging has only recently been applied to the study of cell biological processes in archaea [1], where imaging of halophilic archaea has led to surprising new findings [2-5]. Live imaging of archaea remains challenging, especially when compared to bacteria, because archaea tend to be mechanically soft and several representatives are extremophiles, necessitating the development of novel techniques [1, 2]. Nevertheless, the recent discovery of the *Asgard* superphylum, being more closely related to eukaryotes than any previously discovered organism [6, 7] and the fixed images of the first member of this family to be enriched in lab, *Lokiarchaea* [8], make clear the importance of archaeal cell biology for furthering our understanding of eukaryogenesis.

Among the thermophilic archaea, members of the *Sulfolobales* are one of the most widely used as model organism because they are aerobic, easy to handle and have well-established molecular genetics [9-12]. *Sulfolobus* are also the most intensively studied members of TACK archaea, the superphylum closest to eukaryotes after Asgards [13] and extremely important for evolution studies. Importantly, work on *Sulfolobus* was among the first to reveal striking similarities between the cell biology of eukaryotes and archaea. *Sulfolobus* was shown to have a cell cycle that is similar in structure to the eukaryotic cell cycle, with discrete phases of DNA replication, DNA segregation and division [14]. The *Sulfolobus* genome was shown to contain multiple origins of DNA replication [15, 16] that fire in a coordinated fashion, under the control of proteins that have counterparts in origin firing in eukaryotes, including Cdc6/Orc homologues and the CMG complex [17, 18]. Chromosome organization in *Sulfolobus* is partitioned into transcriptionally active and inactive domains [19], as it is in eukaryotic genomes, and the machinery driving *Sulfolobus* cell division includes homologues of ESCRT-III and VPS4 [20, 21], proteins that execute abscission in many eukaryotic cells. Finally, it has recently been reported that proteasomal degradation is required to initiate ESCRTIII-mediated cell division and to reset the cell cycle in *Sulfolobus*, which resembles the role it plays in triggering mitotic exit in eukaryotes [22].

However, despite the many advantages of using *Sulfolobus* as a model archaeal system, to date, it has not been possible to image *Sulfolobus* grow and divide live. To change this, we designed and built a heated chamber that can be used for live imaging of thermophilic microorganisms on an inverted fluorescent microscope. Using this platform, we call the *“Sulfoscope”*, we have characterized the dynamics of division in wildtype and mutant *Sulfolobus acidocaldarius* cells for the first time using a combination of fluorescent membrane and DNA dyes. In doing so, we identify clear coupling between DNA segregation and cell division in *Sulfolobus* and uncover distinct roles for the two contractile ESCRT-III (CdvB1 and CdvB2) in division itself. This demonstrates how an imaging platform capable of maintaining high temperatures can be used to reveal fundamentally new aspects of cell biology in *Sulfolobus* and thermophilic microbes and reveals new striking differences and parallels between aspects of archaeal and eukaryotic division.

## Results

### High temperature live imaging of fluorescently labelled *Sulfolobus* cells

In order to achieve the stable high temperatures (70°C-80°C) required for live imaging of thermophilic archaea, like *Sulfolobus acidocaldarius*, we initially designed a chamber consisting of two individual heating elements: a heating cap and stage (Figures 1A-E). We based our first-generation design on a commercial imaging chamber (Attofluor®, Invitrogen™) (Figure 1A, bottom part) - using a commercial heating stage to heat the imaging chamber from below. However, this led to large (>10°C) temperature gradients throughout the chamber due to the poor conductivity of the glass coverslip and thermal losses to the surrounding room-temperature air. Furthermore, the open top design led to the rapid loss of media from the chamber through evaporation. This could not be resolved using a simple seal, as this resulted in evaporated water condensing onto the underside of the lid.

**Figure 1.**
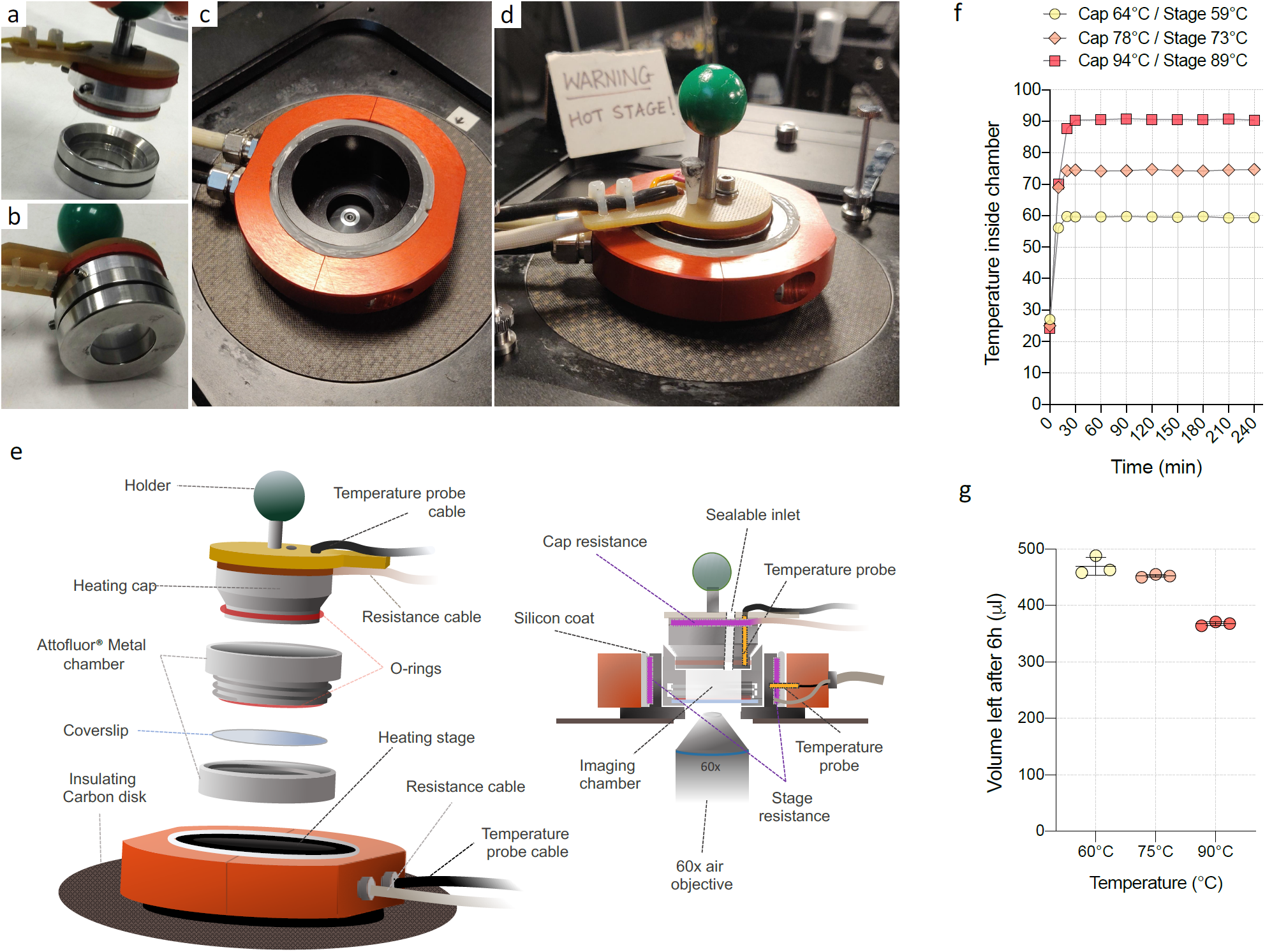
Heating elements for thermophile live imaging. (a) Heating cap and Attofluor^®^ chamber, disconnected. (b) Assembled cap with Attofluor^®^ chamber. (c) Heating stage view from the top. (d) Fully assembled system. (e) Schematics of the heating system. (f) Temperature measurements performed inside the system. Temperature was recorded using a probe inserted inside the imaging chamber, through the sealable inlet. (g) Evaporation measurements after six hours at the desired temperature. Error bars show mean and SD.

To overcome both problems, inspired by PCR machines, we designed and built a heated cap for the chamber (Figure 1A and Figure 1E), which could be sealed using an O-ring (Figure 1B). In order to further improve the temperature control, we built a heated stage (Figure 1C and 1E) that was designed to perfectly accommodate the chamber and cap (Figure 1D). By applying a temperature gradient to the sealed chamber, so that the cap was raised to a temperature 5°C higher than the base, the medium quickly reached a stable equilibrium temperature, that could be maintained for long periods of time with minimal loss of water (Figure 1F, 1G). Detailed illustrations of these chambers can be found in the supplementary material (Supp. Figure 1). Temperatures were measured using a probe that was inserted into the chamber through a small sealable inlet in the cap. Note that this opening also provides a means to inject drugs/media and can be used to introduce microfluidics into the system.

In order to image *Sulfolobus acidocaldarius* cells live using this setup, cells were pre-labelled in solution using dyes (Nile red for membrane and SYBR safe for DNA) that retain their optical properties at high temperature and low pH, and performed better than other tested dyes (e.g. DAPI, Hoechst). Additionally, to successfully extract quantitative data out of time-lapse movies, proper immobilization of *Sulfolobus* cells proved to be a major problem. Unlike bacteria cells, *Sulfolobus* cells appear to be soft and sensitive to mechanical stress (Supp. Figure 2A-B) – as observed by previous groups working with haloarchaea [1, 2]. Although *Sulfolobus* cells could be imaged without immobilization, they had to be kept static for a detailed quantitative analysis of cell division. For this purpose, we employed semi-solid, preheated Gelrite pads (see Material and methods for details). We then identified conditions under which it was possible to combine the two fluorescent dyes, immobilization and two-colour fluorescent light to image *S. acidocaldarius* for a maximum period of 2h. After this, cell divisions became rare. Whereas the membrane dye proved itself to be non-toxic, the use of a DNA dye, as expected and reported for many other cells, affects cell growth (Supp. Figure 2C) and probably accounts in part for the limited time that cells can be imaged (together with potential phototoxicity and the immobilization). Therefore, where possible (e.g. for the study of division symmetry and failures), measurements were performed using Nile Red alone.

### Live imaging reveals the tight coordination of DNA segregation, compaction, and cytokinesis in dividing *Sulfolobus acidocaldarius* cells

Using the *Sulfoscope* we then assessed the dynamics of some of the events that accompany division of the thermophilic archaeon *Sulfolobus acidocaldarius*. In these experiments, *S. acidocaldarius* cells were found to be near perfect spheres, which divide to generate two oval daughter cells (Figure 2A and 2B, Supp. Movie 1). Imaging also revealed a reproducible timeline of events, with coordinated changes in DNA organisation and cell shape (Figure 2A-B and Supp. Movie 1). The first sign that cells were about to divide was a change in the DNA channel 12-16 minutes prior to the first signs of membrane furrowing (Figure 2B and 2D). During this period, the DNA underwent a transition from a very diffuse state to a more compact state in which the nucleoids appeared partially separated (Figure 2B). The onset of furrowing was then accompanied by further compaction of the DNA (measured as a decrease in DNA fluorescence area) (Figure 2D), leading to the formation of two relatively well-defined, spatially separated genomes (Figure 2B, 2D). The completion of cytokinesis in this imaging conditions (immobilized cells with DNA dye) was found to take 10-14 minutes. However, faster division times (around 4 minutes) could be achieved when analysing division in cultures grown non-immobilized and stained with just Nile red (data not shown). Reassuringly, this order of events is extremely similar in movies of dividing *Saccharolobus solfataricus* cells (formerly *Sulfolobus solfataricus* [23]) (Supp. Figure 2D-E and Supp. Movie 2), imaged at 75°C using the same setup, suggesting that the process is conserved between these closely related organisms.

**Figure 2.**
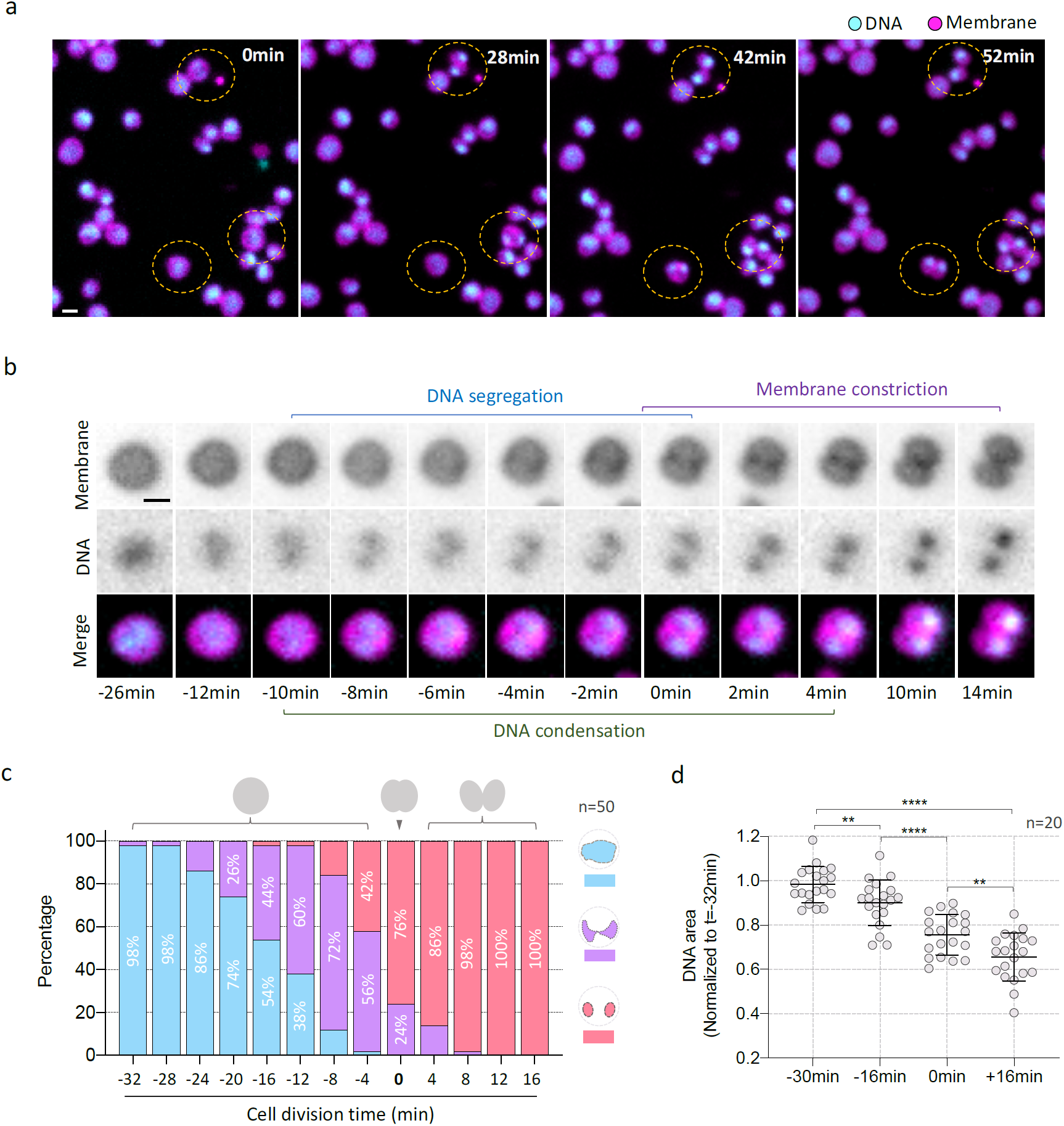
Live cell division of *Sulfolobus acidocaldarius* DSM 639. (a) Time-lapse of a microscopy field showing cells segregating their DNA and dividing. (b) Time-lapse of a selected cell, showing the changes in the membrane and DNA organization trough time. (c) Changes in DNA organization trough time. Cells were separated in three different classes: Diffuse, single disorganized DNA structure (blue); Diffuse, double disorganized DNA structure (Purple); Condensed, well defined DNA structure (Pink). (d) Measurements of DNA area trough time (\*\**P*<0.01, \*\*\*\**P*<0.0001. statistical test: non-parametric Mann–Whitney). Scale bars:1μm. Error bars show mean and SD.

### The role of the non-essential ESCRTIII proteins, CdvB1 and CdvB2, in *S. acidocaldarius* cell division

Risa and collaborators [22], recently put forward a model of ESCRTIII-mediated division for *S. acidocaldarius* based on data acquired from using fixed cells. These data suggested that cell division is regulated as follows: a rigid CdvB division ring is formed at the equatorial plane of dividing *Sulfolobus* cells, which functions as a template for the assembly of a contractile ESCRTIII heteropolymer based on two other ESCRTIII proteins, CdvB1 and CdvB2. Sudden degradation of CdvB via the archaeal proteasome then triggers division by allowing the CdvB1 and CdvB2 ring to undergo a change in curvature to induce membrane constriction and scission [22]. Under this model, CdvB1 and CdvB2 are treated as being equivalent in terms of their ability to drive scission [22]. This fits with their co-localisation [22] and is in keeping with previous studies that identified CdvB1 and CdvB2 as close paralogues that share 65% amino-acid identity and 80% similarity. We wondered, though, if live-cell inspection of cell division events using the *Sulfoscope* could reveal more details about the roles of CdvB1 and CdvB2. To test this, we generated in-frame deletion mutants of *cdvB1* and *cdvB2* in *S. acidocaldarius* background strain MW001 (Figure 3A and 3B) and imaged these cells live (Supp. Movie 3). Interestingly, while growth was severely impaired in the *cdvB2* deletion strain, the deletion of CdvB1 only had a marginal impact on growth (Figure 3A).

**Figure 3.**
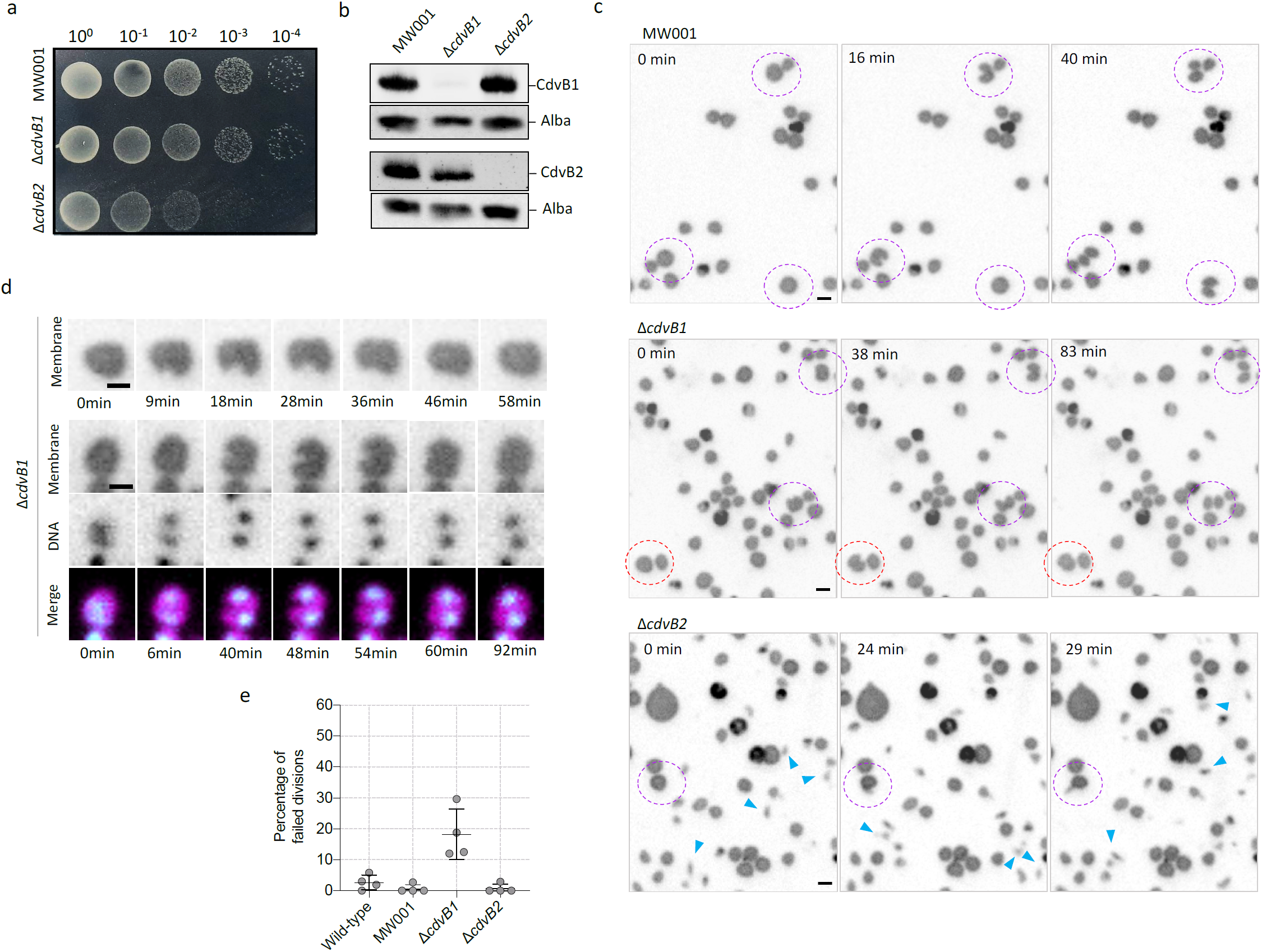
Effects of deletion of *cdvB1* and *cdvB2* in cell division and cell growth in *Sulfolobus acidocaldarius.* (a) Growth of MW001 (background strain), Δ*cdvB1* and Δ*cdvB2* in BNS plates, showing that growth of Δ*cdvB2* is greatly affected. (b) Western blots of MW001, Δ*cdvB1* and Δ*cdvB2* using CdvB1 and CdvB2 antibodies. As a loading control, we used the DNA binding protein Alba. (c) Time lapse microscopy showing fields of MW001, Δ*cdvB1* and Δ*cdvB2.* Successful divisions are shown in purple dashed circles, whereas failed divisions are shown in red dashed circles. The blue arrowheads point to small cells that constantly move in the imaging field. (d) Time-lapse of selected cells of Δ*cdvB1* that fails division, resulting in the fusion of the daughter cells, following initial membrane furrowing. (e) Quantification of the frequency of division failures in the Wild-type, MW001, Δ*cdvB1* and Δ*cdvB2.* Four independent movies were used to evaluate the failure frequency, and at least 100 divisions in total were considered for each strain. Scale bars: 1μm. Error bars show mean and SD.

While Δ*cdvB1* cells showed similar size compared to the background strain, MW001 (Figure 3C and Supp. Movie 3), deletion of CdvB1 led to some cells to fail midway during division (Figure 3D-E and Supp. Movie 4). In such cases, a sudden halt in cytokinesis caused the nascent daughter of Δ*cdvB1* cells to fuse back together to generate single cells that carry both copies of the spatially separated and compact chromosomes (Figure 3D). Such failures, in our imaging conditions, account for 18% of the Δ*cdvB1* divisions, an event that was rare (1-2%) in the other strains (Figure 3F). Thus, while non-essential, CdvB1 is required to ensure that division is fail-safe in *Sulfolobus acidocaldarius*.

However, as suggested by the colony phenotype, the impact of loss of CdvB2 on division was much more profound. At a first glance, a significant cell size variation can be observed (when compared to MW001), as well several small, irregularly shaped cells that are not properly immobilized by the pad (Figure 3C and Supp. Movie 3). Strikingly, a closer look revealed that the small cells in the culture are generated by asymmetric divisions. Moreover, the asymmetry of the divisions in Δ*cdvB2* cells was very variable (Figure 4A-D, Supp. Movie 5). In most cases, each of the two daughter cells generated by an asymmetric division retained one of the two separated chromosomes (Figure 4B). However, in some cases, extreme mispositioning of the furrow led to the formation of ghost cells (Figure 4C), generating a subpopulation of large cells with extra chromosomes (Supp. Figure 3). This was also visible as a high variance in the size of newly born G1 cells (and to a lesser extent G2 cells (Supp. Figure 4)) – as indicated by the analysis of populations of fixed cells labelled with a DNA dye by flow cytometry (Figure 4E and 4F). Importantly, the division asymmetries and the cell size variation seen in the Δ*cdvB2* strain could be rescued by the ectopic expression of CdvB2 from an inducible promoter (Figure 4D-F, Supp. Figure 4, Supp. Movie 6), showing that these defects are due to a lack of CdvB2.

**Figure 4.**
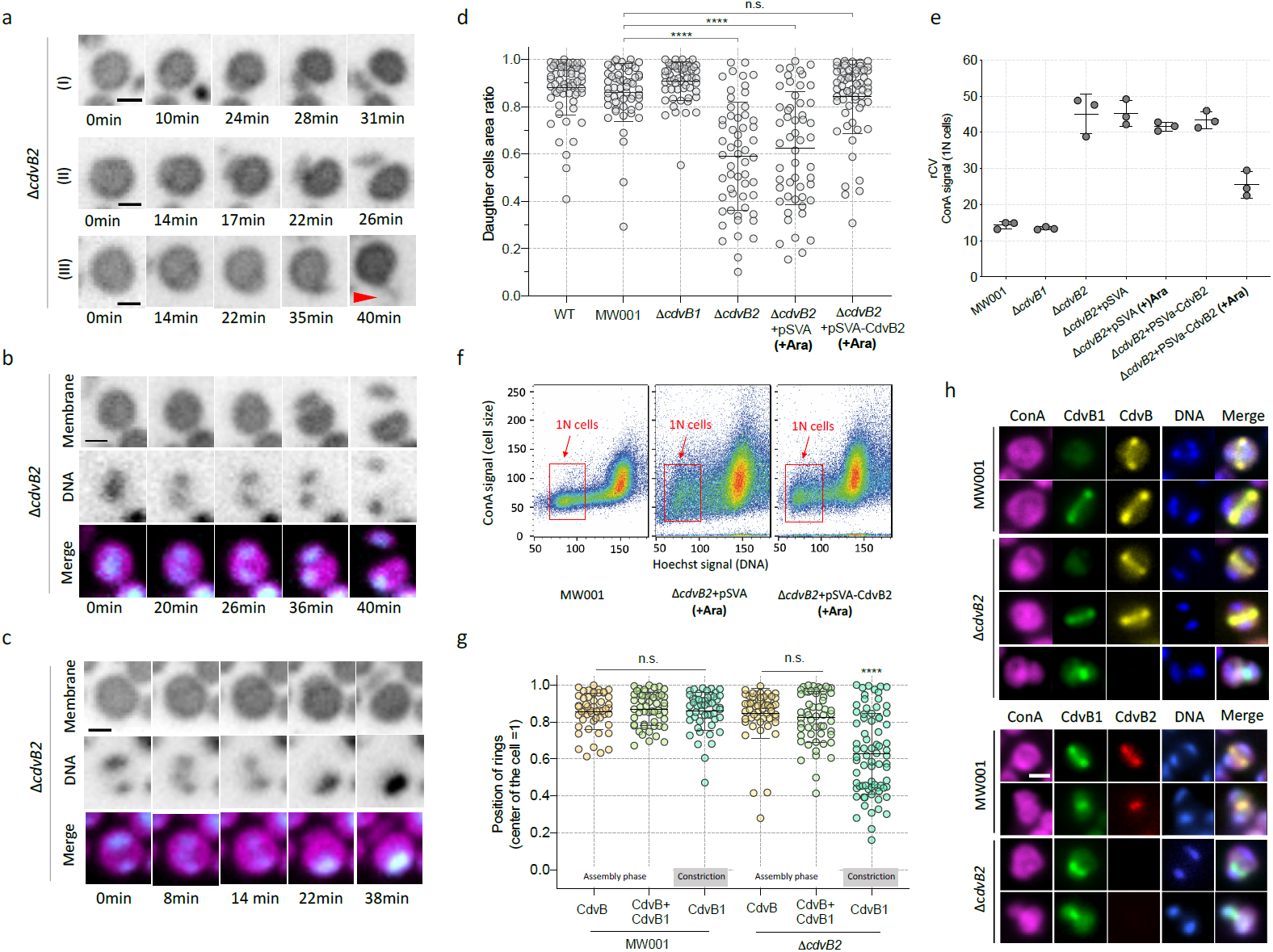
Cell division is greatly disrupted in the absence of the ESCRT-III protein CdvB2. (a) Asymmetric divisions, from moderate (I), to strong asymmetry (II and III). Red arrowhead shows a very asymmetric daughter cell. b) Asymmetric division, with both cells ending up with DNA inside (c) Asymmetric division of a cell in which a ghost cell is formed (d) Daughter cell area ratios of the tested strains. The value of 1 represents a perfect symmetric division, whereas smaller values indicate increasing asymmetry. Divisions from of at least three independent movies were used, and at least 50 cells were evaluated for each strain (\*\*\*\**P*<0.0001, n.s.: not significant. statistical test: non-parametric Mann–Whitney). (e) Size variation of newly divided cells in ethanol fixed cells, estimated by Flow-cytometry. The plot shows Robust coefficient of variance (rCV = st.dev/median), considering ConA-alexa 647 signal, that binds to cell envelope (f) Flow-cytometry charts, showing size variation of MW001, Δ*cdvB2*+pSvA (Induced) and Δ*cdvB2+*pSvA-CdvB2 (Induced), showing that the expression of CdvB2 in a plasmid partially rescues the phenotype. 1N cells indicates cells with one copy of the chromosome (recently divided). (g) ESCRT-III division rings positioning quantification, in MW001 and Δ*cdvB2* ethanol fixed cells and immunostained with CdvB and CdvB1 antibodies. Rings positioning were estimated accordingly to the order of events of division, described previously (Risa et al., 2019) (\*\*\*\**P*<0.0001, n.s.: not significant. statistical test: non-parametric Mann–Whitney). (h) Immunofluorescence of MW001 and Δ*cdvB2* using anti-CdvB, CdvB1 and CdvB2. Scale bars: 1μm. Error bars show mean and SD.

To determine the precise timing of the error in the positioning of the furrow relative to the DNA, we had to revert to immunostaining of fixed cells, given the current absence of thermophilic fluorescent protein that can be used in *Sulfolobus.* For this, we immunostained fixed Δ*cdvB2* cells for the ESCRT-III proteins (CdvB, CdvB1 and CdvB2). Using the sequence of events described previously [22], we were able to infer the position of the ESCRT-III rings at different stages in the division process in these samples based upon the presence or absence of CdvB (Figure 4G). CdvB was found correctly positioned in the centre of MW001 and Δ*cdvB2* (Figure 4H). The same was true for cells that co-stained for CdvB and CdvB1, with a few exceptions for mispositioned rings in Δ*cdvB2* cells (Figure 4G). Therefore, no obvious differences were observed during the assembly phase. However, in Δ*cdvB2* cells starting to divide (constriction phase), as the result of proteasome-mediated degradation of CdvB [22], CdvB1 was found asymmetrically positioned, while it remained correctly positioned at the cell centre in the MW001 control (Figure 4G, 4H). These data suggested that CdvB2 is recruited (along with CdvB1) to the correct position at the cell centre by the CdvB ring, where it is required to stop the CdvB1 ring slipping relative to the chromosomes after the degradation of CdvB. This slipping was also visible as a gradual slippage of the nascent furrow, visible as a concentration of Nile red, in some *cdvB2*Δ cells undergoing asymmetric divisions (compare Figures 2B and 4A).

## Discussion

Here we describe the development of the “*Sulfoscope”*, an imaging platform that makes it possible to image *Sulfolobus* cell divisions live. Our first use of the *Sulfoscope* revealed a tight coupling between DNA compaction, nucleoid separation and membrane deformation during division (Figure 2D). Strikingly, these events appear to occur in a defined order in the wildtype. This begins with the genome losing its relatively diffuse organisation as it separates to form two relatively compacted chromosomes (Figure 2C, 2D). This is followed by a relatively sudden increase in the rate of DNA compaction (over the course of a few frames), which coincides with the onset of furrowing - an event that appears to pull the genome out of the way as the division ring starts to narrow. Interestingly, the same sequence of events was observed for *S. solfataricus* (Supp. Figure 2), suggesting that it is likely conserved across *Sulfolobus* species. Thus, in *Sulfolobus*, DNA compaction may help to clear the DNA out of the way of the furrow as it closes.

By combining molecular genetics and live cell imaging, we were also able to use these data and the *Sulfoscope* to define homologue-specific roles for the two different ESCRT-III proteins that form part of the contractile division ring [22]. This is important, because these proteins appear to play an analogous role in the membrane remodelling events that accompany cell division in *Sulfolobus* to that played by ESCRT-III homologues in cell abscission in eukaryotes, a process that is still not totally understood [24]. While CdvB1 and CdvB2 are co-localised during division (Figure 4H) and are close homologues, which were likely generated by a relatively recent gene duplication within the family, this analysis revealed that they perform distinct functions to ensure that division is robust and symmetric. Thus, cells that lack CdvB1 undergo occasional cytokinesis failure, resulting in the fusion of daughter cells, as occurs in ESCRTIII-mutants in some eukaryotes [25, 26]. By contrast, the loss of CdvB2 appears to compromise the stable positioning of the ring as it constricts – so that CdvB1 rings slip in the mutant causing profound defects in division symmetry (Figure 4D). As a consequence of such slippage, during the constriction phase, a proportion of divisions generate ghost cells and daughter cells with twice the normal DNA complement (Figure 4C and Supp. Figure 3). Thus, while it is possible that the two proteins might function as a heteropolymer in the wildtype, as has been shown for many eukaryotic ESCRT-III homologues [27, 28], they appear to perform distinct but complementary functions in ring contraction. Strikingly, these data also show that cells can generate the forces required to deform the membrane to divide using just one of the two contractile ESCRT-III rings. These data also show, unexpectedly [21], that both CdvB1 and CdvB2 can be recruited by the CdvB ring in *S. acidocaldarius.*

Signs for such distinct functions for CdvB1 and CdvB2 can be found in the literature. For example, deletion of *cdvB2* rendered cells slower-growing than did the deletion of *cdvB1* in a previous study [29], but no specific phenotype besides growth could be addressed for such mutants. Also, whereas *cdvB1* deletion cells could be generated in *Sulfolobus islandicus* [9], *cdvB2* was found to be an essential gene in this organism. In addition, Liu and collaborators [30] observed distinct effects of overexpressing truncated versions of CdvB1 and CdvB2 in *S. islandicus* by imaging static cells at room temperature. In their study, the over-expression of the truncated versions of CdvB2 version generated cells with defects in the final stages of abscission [30] and protrusions similar to midbodies, whereas the overexpression of truncated CdvB1 generated a population of connected cells, similar to a chain. While we were unable to observe such phenotypes in our analysis, determining the precise roles of individual proteins in a dynamic process like division greatly benefits from live cell imaging, something that could not be achieved before. It is only by imaging loss of function mutant cells live as they divide, in strains where we can switch the expression of a rescue construct on or off, that the roles of these proteins in ensuring a robust division becomes clear.

These findings also have parallels in eukaryotes. In mammalian cells, for example, the two ESCRT-III proteins CHMP2A and CHMP2B, which we previously assumed to be functional homologues, were recently found to contribute differently to membrane remodelling and to have different affinities for the membrane [31]. Moreover, in *in vitro* studies, different eukaryotic ESCRT-III proteins have been shown to work together to increase membrane binding of the heteropolymer [32]. Our data suggest that the same might apply in *Sulfolobus*, where two different ESCRT-III proteins [21, 30], CdvB1 and CdvB2, act cooperatively to ensure proper membrane binding and constriction - preventing ring slippage or cytokinesis failure. We think this work also makes the case for *Sulfolobus* being a simple, well-defined system in which to study ESCRTIII-dependent division. Thus, in the future we aim to use a combination of live imaging, CryoEM, in vitro assays and modelling [22] to determine how ESCRT-III proteins are able to work as single or heteropolymers to apply mechanical forces required to deform a membrane and induce scission – something that remains incompletely understood in eukaryotes [24].

Finally, while we have used the *Sulfoscope* to reveal fundamental novel aspects of cell division, we anticipate that this high-temperature imaging platform can be applied to shed light on other exciting areas of cell biology in *Sulfolobus*, including DNA remodelling [19, 33], swimming [34], conjugation [35, 36], viral infection [30, 37], competition [38, 39] and cell-cell fusion [40]. Live imaging of *Sulfolobus* will also uncover entirely new cell biology in these organisms, since so far, we have not been able to watch them alive. In addition, it will also be relevant for the study of other thermophilic microbes, from eukaryotes to bacteria [41, 42]. Therefore, we hope that our work can open new avenues of research and leading further developments in the field of live imaging of thermophiles and *Sulfolobus*.

## ACKNOWLEDGEMENTS

The authors would like to thank the Baum lab for their input throughout the project. We would like to thank Juan Manuel Garcia for help early on in this project. Tobias Härtel, Pedro Matos Pereira and Alexandre Bisson for discussions about fluorescent probes; Alexander Wagner for providing the plasmid pSVAaraFX-stop. Alexandre Bisson and Jan Löwe for comments on the manuscript. Mateusz Trylinski for helping with movies. AAP was supported by HFSP LT001027/2019. fellowship. DRM was supported by the BBSRC (BB/K009001/1). GTR was supported by the MRC PhD studentship award (MC_CF12266). GD was funded by a European Union Marie Sklodowska-Curie Individual Fellowship (704281-CCDSA). In addition, AAP, SC and GD all received support from the Wellcome Trust (203276/Z/16/Z). Research was supported by Wellcome Trust (203276/Z/16/Z) and VW life? Grant (Az 96727) with additional support for BB and RH provided by UCL.

## AUTHORS CONTRIBUTION

AAP, DRM, GD, RH and BB conceived the study. Microscope design and construction and commercial heating stage testing was carried out by AAP, SC, GD, DRM and RH. Design of the heating cap and heating stage was carried out by AAP, JR, CR and MR. JR and MR constructed the heating cap and heating stage. AAP and DRM established the imaging methodology and dyes used. Live-imaging and imaging analysis were performed by AAP and DRM with help from SC. Genetics and strains were performed by AAP, MVW and KNS. Immunofluorescence and Western-Blots were performed by AAP. Flow Cytometry were performed by GTR. Chamber measurements were performed by AAP. The bulk of data analysis was carried out by AAP, and the figures and text were prepared by AAP and BB, with input from all other authors.

## Material and Methods

### Cap/stage construction and development

The heating system was designed to be combined with the Attofluor^®^ chamber. The metal elements in the cap and the heating elements were built with aircraft aluminium (AL 7075 for the cap and stage and AL 2024 for the metal collars) using a lathe to obtain the final shape. Screws used are stainless steel 304. The insulating disk was built with carbon fibre. The top part of the heating cap was built using fibreglass and a silicon rubber, which separates the metal and the fibreglass and holds the electric resistance (thin film resistance). The temperature of the cap and the chamber were controlled by two independent heating systems attached to the heating elements. The temperature used in the cap is a thermocouple type T; the one used for the chamber is a miniPT100. Detailed information about the design of the chamber is shown in Supp. Figure 1.

### Chamber measurements

For measurements of the temperature inside the chamber, a digital thermometer with a probe (Signstek 6802 II with a 2k type probe) was used. The probe was inserted inside the chamber trough the sealable inlet (Figure 1) and positioned exactly at the middle of the chamber, touching the glass. 500μL of Ultrapure water was added to the chamber, since in our imaging conditions, liquid media is always present. Readings were taken initially every 10 minutes (for the first 30 minutes) then followed up for 30 minutes. A closer follow up was performed as well, following the temperature continuously for half an hour for every tested temperature. We never observed a temperature variation greater than 0.5°C during the whole duration of the measurements. For the evaporation assays, 500μL of BNS media was added to the chamber, and the weight of the chamber was measured using an analytical scale. The pre-heated system was them closed. After six hours, the heating system was turned off. After cooling back to room temperature, the Attofluor chamber was removed and the weight was recorded again, and the difference in the final weight was considered as loss of liquid due to evaporation.

### Strains constructions and growth conditions

The *S. acidocaldarius* wild-type (DSM639), *S. solfataricus* wild-type (P2) Δ*cdvB2*+pSVAaraFX-stop (empty plasmid control) and Δ*cdvB2*+pSVA-CdvB2 were grown at 75°C in Brock Salts medium supplemented with 0.1% NZ-amine and 0.2% Sucrose, final pH corrected to 3.5 with H_2_SO_4._ Uracil was added to the final concentration of 20μg/mL when working with uracil auxotrophic strains MW001, Δ*cdvB1* and Δ*cdvB2* strains. The density of liquid cell cultures was maintained at OD_600nm_ values between 0.05 and 0.3 as measured by a spectrophotometer. Table of strains used in this work, as well plasmids, can be found in the supplementary material (Table 1 and Supp. Table 1). Deletion of genes was performed using the “pSVA431 method” as previously described by Wagner and collaborators [10].

**Table 1.** Stains and plasmids used in this work.

| Strain | Background | Description | Source |
| --- | --- | --- | --- |
| *S. solfataricus* P2 | Wild-type | Wild-type | DSMZ |
| *S. acidocaldarius* DSM639 | Wild-type | Wild-type | DSMZ |
| MW001 | DSM639 | Uracil auxotrophic background strain $\Delta pyrE$ ($\Delta bp$ 91–412) | Wagner *et al.*, 2012 |
| $\Delta cdvB1$ | MW001 | $\Delta cdvB1$ (Saci_0451) | This work |
| $\Delta cdvB2$ | MW001 | $\Delta cdvB2$ (Saci_1416) | This work |
| $\Delta cdvB2$+pSVA | $\Delta cdvB2$ | $\Delta cdvB2$ transformed with pSVA empty plasmid. | This work |
| $\Delta cdvB2$+pSVA-CdvB2 | $\Delta cdvB2$ | $\Delta cdvB2$ transformed with pSVA containing the wild-type gene of CdvB2, under an inducible promoter controlled by Arabinose. | This work |

| Plasmid | Backbone | Description | Source |
| --- | --- | --- | --- |
| pSVA431 | - | Backbone of deletion plasmid containing a *pyrEF* and *LacS* cassette from *S. solfataricus* | Wagner *et al.*, 2012 |
| pSVA1895 | pSVA431 | Used for deletion of *cdvB1* (Saci_0451) | This work |
| pSVA1896 | pSVA431 | Used for deletion of *cdvB2* (Saci_1416) | This work |
| pSVAaraFX-stop | - | Backbone of expression plasmid harbouring an arabinose inducible promoter containing a *pyrEF* cassette from *S. solfataricus* | Alexander Wagner (unpublished) |
| pSVAaraCdvB2 | pSVAaraFX-stop | Used for expression of CdvB2 (Saci_1416) | This work |

For the Δ*cdvB2*+pSVAaraFX-stop and Δ*cdvB2*+pSVA-CdvB2 rescue experiments, expression was induced by adding 0.15% w/v (final concentration) of L-arabinose to freshly diluted cultures. Cells were grown overnight with L-arabinose and collected the next morning for live-imaging, immunofluorescence and flow cytometry, at an OD_600nm_ of 0.1 to 0.25. The experiment was repeated three times, on independent days.

### Live imaging methodology

All the manipulation of cells and materials prior to live imaging at high temperature were conducted in a way to minimize temperature loss during the transport from the incubators to the microscope. In order to achieve this, metal beads were placed inside a container that was heated overnight at 75°C. Aliquots of the cultures (5mL to 10mL) were transferred to previously clean and heated flasks placed inside the metal beads container, and then transported to the microscope. When done properly, cells suffer from minimal heat stress, and cell division can be imaged immediately after the start of the image acquisitions. The commercial Attofluor chamber and coverslips were washed thoroughly with BNS media, followed by a wash with ultrapure water before use. We noticed that unclean coverslips can potentially damage *Sulfolobus* membrane. The use of Nile red membrane allowed us to detect if cells were damaged prior to cell imaging (Supp. Figure 2). The assembled chamber containing the clean coverslip needs to be pre-heated before imaging starts. This can be achieved during the washing/drying steps.

For live imaging of *S. acidocaldarius*, Nile red (Sigma) was used at 2.5μg/mL final concentration and SYBR safe (Invitrogen), 10,000x concentrated was used at a final dilution of 0.5x to 1x. Cells were imaged at an OD_600nm_ of 0.1 to 0.35. For imaging immobilized cells, we used heated semi-solid BNS pads (0.6% Gelrite, 50% non-pH BNS, with 20mM final concentration of CaCl_2_). Pads were prepared by cutting circular (8mm) disks from a semi-solid BNS media petri dish, prepared as described above (25mL per plate). To facilitate the handling, a 13mm circular coverslip was used to support the pad after it was removed from the plate. The pads, now inside a closed petri dish, were placed at an incubator at 75°C degrees for 20 min prior to imaging and transported inside heated metal beads container (as described above) to the microscope. After loading 400μl of the cell culture inside the chamber, the heated pads were placed on top of the cells (using forceps, holding the 13mm coverslip with the pad on top). The chamber was quickly closed and positioned for imaging. We observed that heating up the pads prior to imaging not only prevented heat shock, but also slightly dried the pads. This causes the cells to accumulate at the border of the pads once placed inside the chamber. Movies were therefore acquired at those regions, since more cells can be found there and also those regions allow some degree of cell movement as well. Areas in which cell membrane is clearly stressed (Supp. Figure 2) were avoided when imaging with pads. Imaging immobilized cells, with DNA dye, Membrane dye and constant exposure to light is limited to two hours, probably to the combined toxic effect of the DNA dyes and light exposure. Nevertheless, even without the use of dyes, additional issues arise when imaging cells over longer periods, including water loss and insufficient oxygenation, both issues that can be overcome with the use of microfluidics.

### Imaging acquisitions, cell measurements and quantifications

Images were acquired using a Nikon Eclipse Ti-2 inverted microscope, equipped with a with a Prime 95B sCMOS camera (Photometrics). A 60x air objective (Plan Apo 60x/0.95 objective, Nikon) combined with the 1.5x additional magnification from the Ti-2 microscope body was used, resulting in an effective magnification of 90x (Pixel size = 0.1235 μm). For membrane imaging, Nile red was illuminated with 550nm LED illumination at 20% of maximum intensity, while for DNA imaging SYBR safe was illuminated with 470nm LED at 25% of the maximum intensity (pE-4000 LED illuminator, CoolLED). Exposure time was limited to about 40 milliseconds for each channel. Whenever DNA was imaged together with Nile red, images were acquired every 2 minutes. For measurement of division asymmetry and failure division events, only Nile red was used, and images were acquired every 30 seconds to every 1 minute. For all live imaging analysis here presented, cells from at least 3 different movies, performed on independent days, were used. Cells and DNA area were measured using Fiji (ImageJ). For correction of XY axis drifting in the images, the ImageJ plugin StackReg (http://bigwww.epfl.ch/thevenaz/stackreg/) was used [43]. Focal drift was avoided by using the inbuilt Perfect Focus System of the microscope body.

### Immunofluorescence and flow cytometry

Immunofluorescence, Flow cytometry methodologies and antibodies description used in this work are reported in Risa et al., (2019). Cells were fixed sequentially in ice-cold ethanol until reaching the final concentration of 70% EtOH. For the analysis, cells were centrifuged (3min at 8,000xg) and resuspended in PBSTA (PBS with Tween20 (0.2% v/v final concentration) + BSA (1% w/v final concentration) for 5 min for rehydration. After rehydration, cells were incubated with 100μl of PBSTA with primary antibodies for 2 hours, at 500rpm, room temperature, in a small thermomixer. Cells were washed twice in PBSTA and them incubated in 100ul of PBSTA with secondary antibodies, mixed with Concavalin A conjugated with 647 alexa fluor at a 200μg/mL final concentration for 1 hour and incubated at 500rpm, room temperature. Cells were washed twice again, resuspended in PBSTA and Hoechst was added to the final concentration of 1μg/mL, to visualize DNA. Cells were them imaged or analysed by flow cytometry, as described in Risa et al., 2019.

### Western blotting

*S. acidocaldarius* pellets were lysed in 100-250μl lysis buffer (TK150 buffer, supplemented with DNAse I, and EDTA-free protease inhibitor cocktail, 0.1% Triton X-100). Cells were disrupted in a sonicator (Diagenode) and subsequently centrifuged at 4°C (10,000 ×g for 15 min). Supernatant was collected and protein concentration was measured using Bradford reagent (Bio-Rad). About 8-15μg of total protein extract was resolved on NuPAGE 4-12% Bis-Tris gels (Invitrogen) using MES SDS running buffer (Invitrogen). Proteins were transferred to a 0.45μm nitrocellulose membrane. Membranes were blocked with PBST (PBS, 0.2% Tween20) with 5% non-fat milk. Primary and secondary antibodies were diluted in this same buffer. Membranes were incubated with primary antibodies overnight at 4°C. Primary antibodies anti-CdvB1, anti-CdvB2 and anti-Alba were used as described in Risa, 2019. Next, the membranes were washed with PBST and incubated with Li-COR IRDye 680CW or 800CW secondary antibodies diluted 1:10000 for 1 hour at room temperature. Finally, membranes were washed with PBST and developed using Li-COR Odyssey Infrared Imaging System. Images were analysed on ImageJ.

