## Supplemental Data 1 for "Live cell imaging of the hyperthermophilic archaeon *Sulfolobus acidocaldarius* identifies complementary roles for two ESCRTIII homologues in ensuring a robust and symmetric cell division"

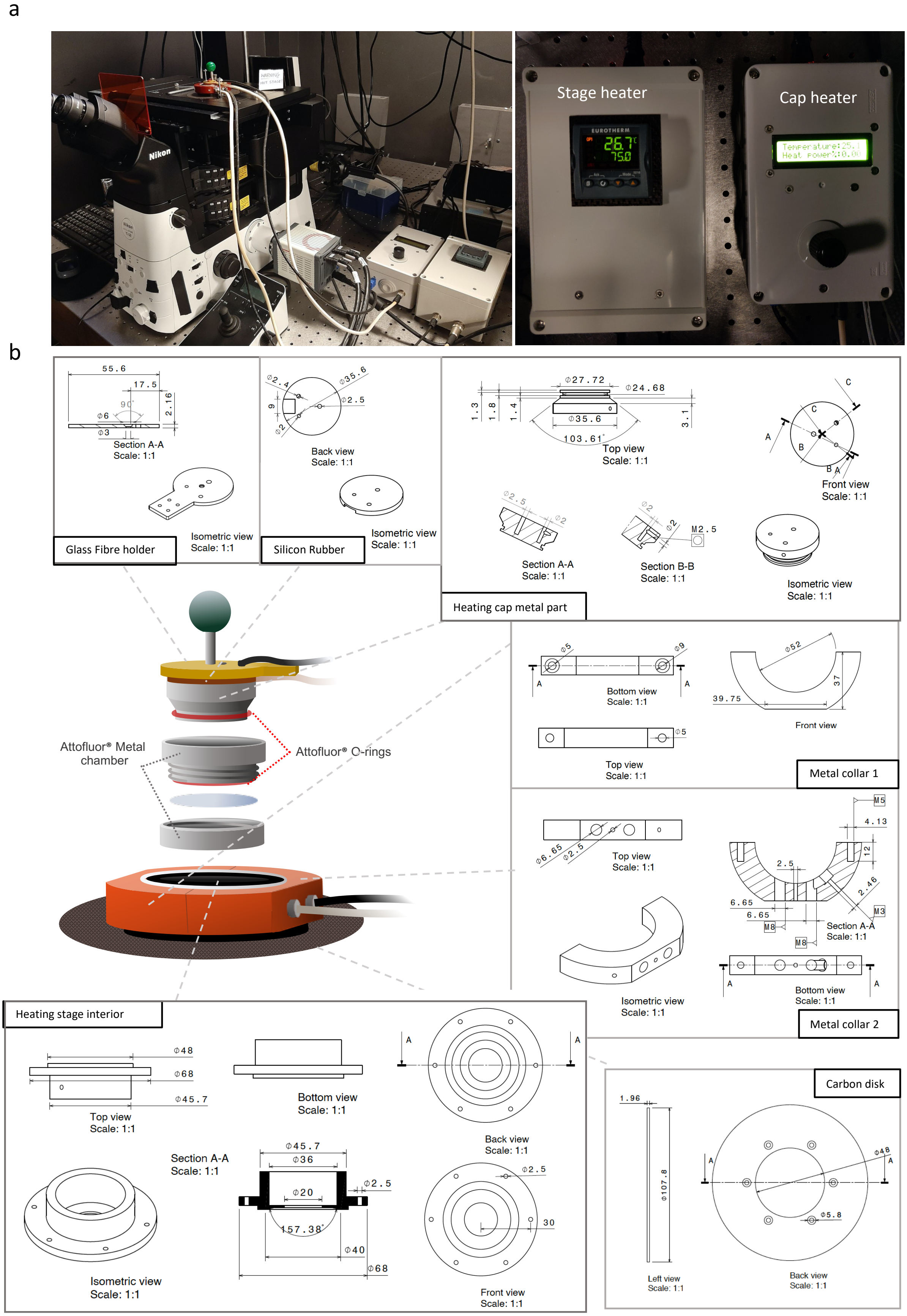


**Supp. Figure 1.** Heating stage and heating system. (a) Picture of the assembled heating system in the microscope, with the heating elements and the respective control systems (shown in the right panel). (b) Schematics of the heating elements, with more detailed information about dimensions and individual parts.

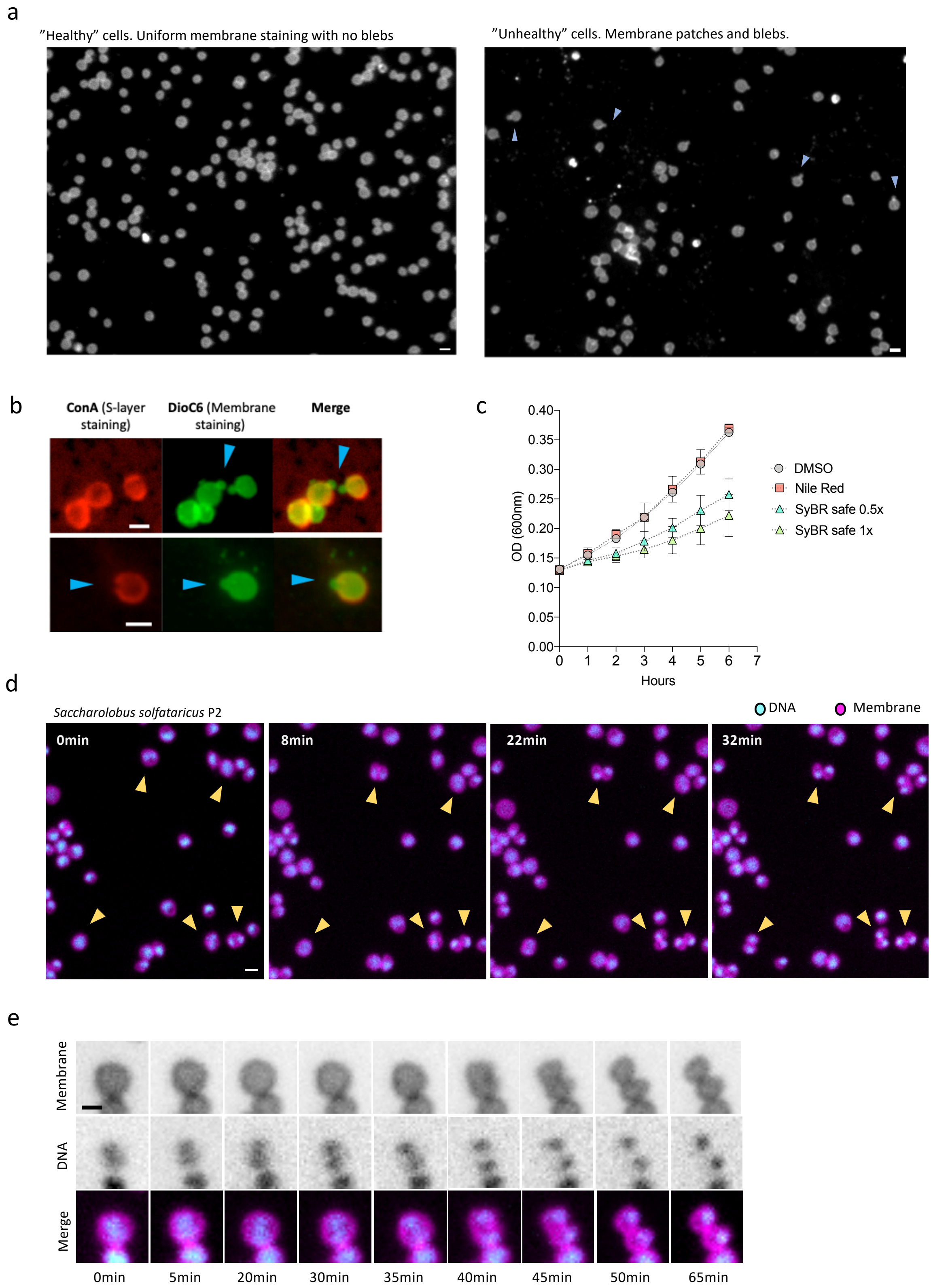


**Supp. Figure 2.** Mechanical stress and unclean coverslips can damage the membrane of *S. acidocaldarius*. (a) Image of *S. acidocaldarius* cells at 75°C illustrating the differences between healthy cells and stressed cells. Stressed cells have typical blebs and membrane patches. For the live imaging, areas like these were avoided. (b) Images of *S. acidocaldarius* (imaged at room temperature) stained for its membrane and S-layer, illustrating that the blebs formed are in fact envelope stress, since those are not coated by S-layer (c) Growth curve of *S. acidocaldarius* treated with Nile Red, SYBR safe and control, at the concentrations used at this work. Error bars show mean and SD. (d) Live imaging of *Saccharolobus solfataricus* P2 (Former *Sulfolobus solfataricus* P2). Time-lapse of a field of view, showing 5 dividing *S. solfataricus* cells. (e) Time-lapse of a selected *S. solfataricus* cell, showing the dynamics of DNA organization and cell division. Scale bars: 1μm

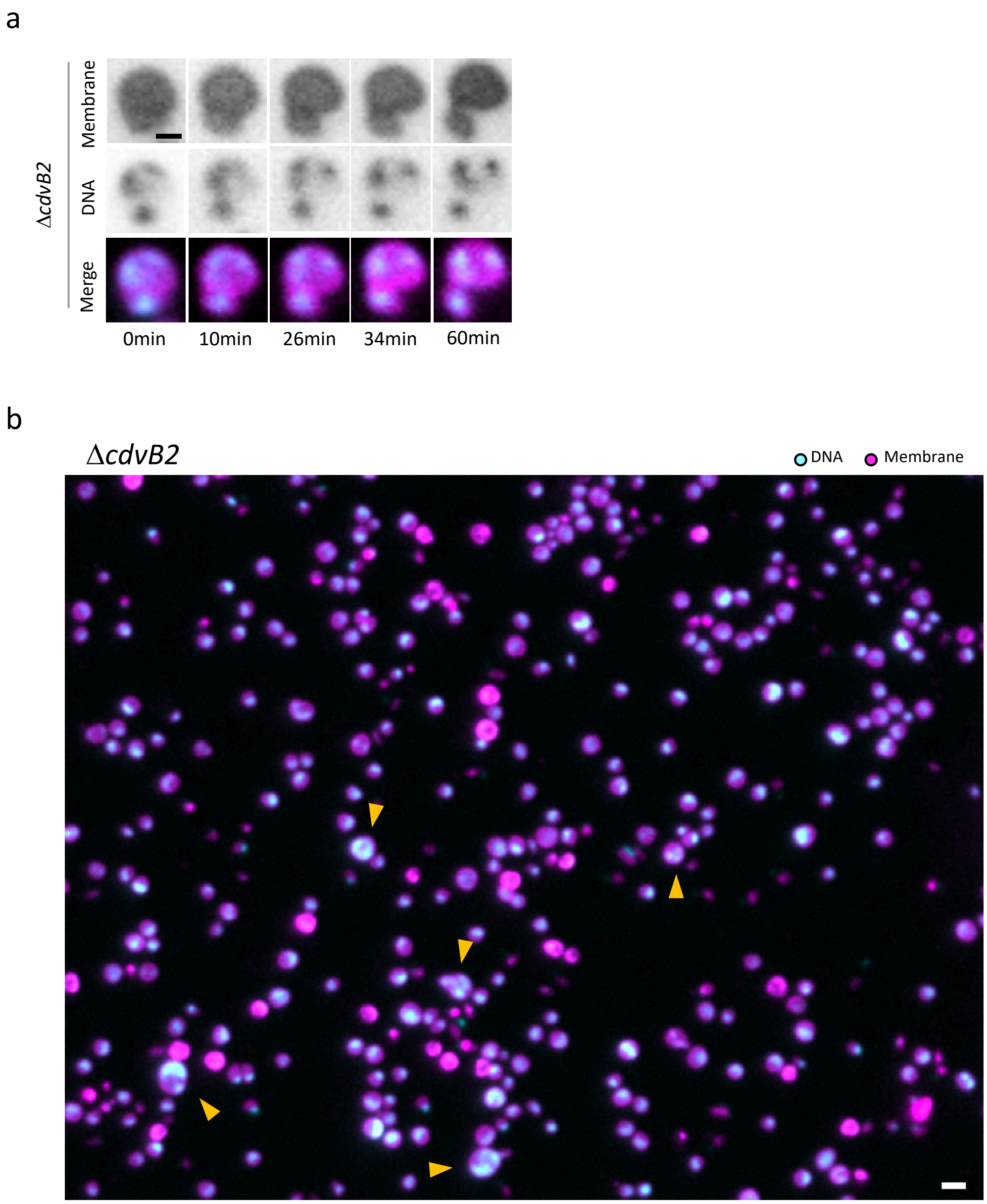


**Supp. Figure 3.** (a) Example of cell division in a large, 2N+ cell. (b) Field of view showing examples of large cells (yellow arrows) with extra copies of chromosomes in the Δ*cdvB2* population. White scale bar: 2μm. Black scale bar: 1μm

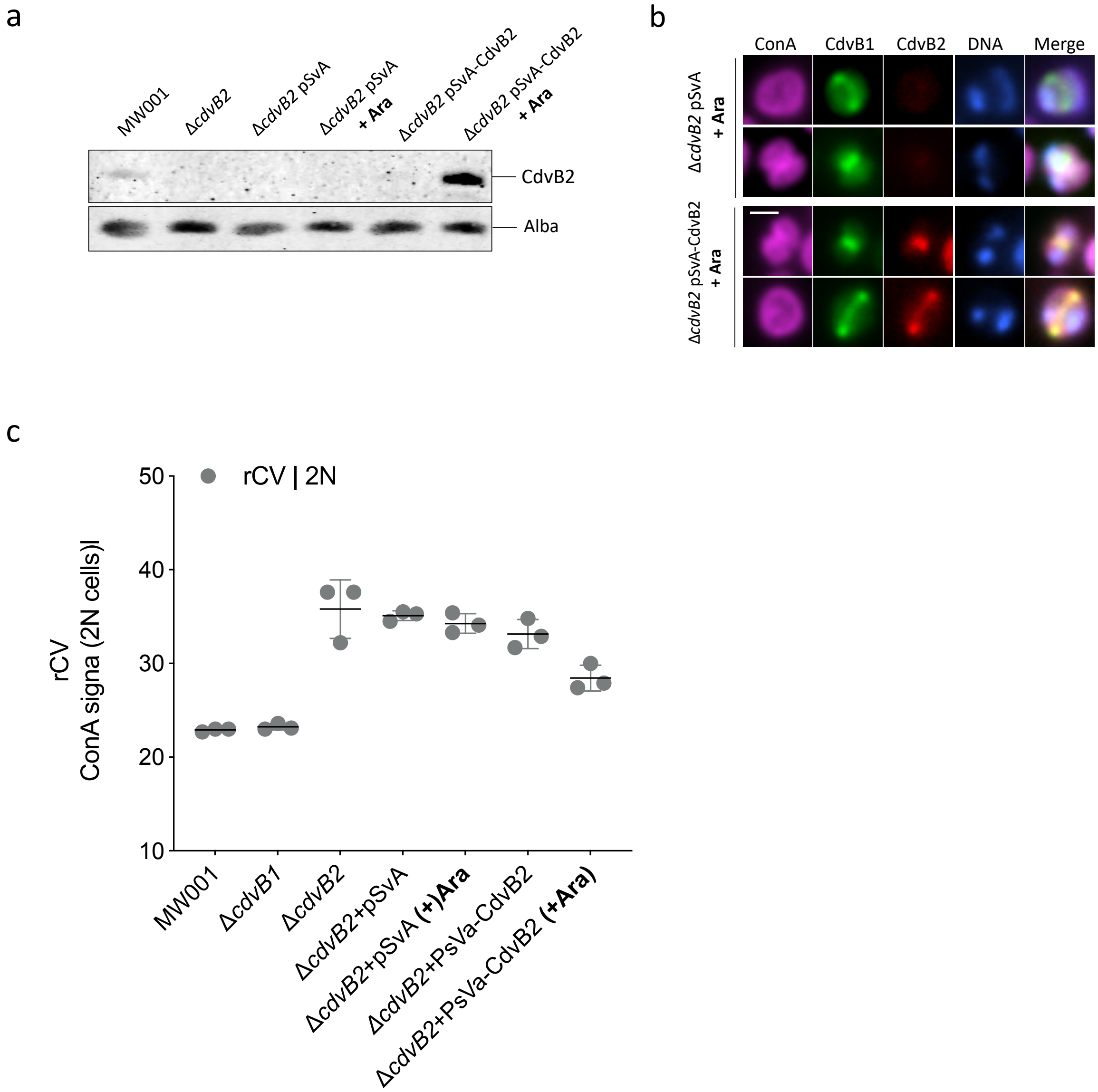


**Supp. Figure 4.** Rescue experiments of the Δ*cdvB2* strain. (a) Western-blot against CdvB2 and Alba proteins, showing the expression of CdvB2 in a plasmid after the addition of the inducer L-arabinose. (b) Immunostaining of the Δ*cdvB2+*pSVA-CdvB2 and Δ*cdvB2+*pSVA (empty plasmid), showing that after the induction CdvB2, the formation of CdvB2 rings is restored. (c) 2N cell size variation, estimated by flow cytometry in fixed cells. rCV = Robust coefficient of variance. Error bars show mean and SD
